# Amount of Pannexin 1 in smooth muscle cells regulates sympathetic nerve induced vasoconstriction

**DOI:** 10.1101/2022.09.07.506995

**Authors:** Luke S. Dunaway, Marie Billaud, Edgar Macal, Miranda E. Good, Christopher B. Medina, Ulrike Lorenz, Kodi Ravichandran, Michael Koval, Brant E Isakson

**Author notes:** to whom correspondence should be addressed* PO Box 801394, University of Virginia School of Medicine, Charlottesville, VA 22908 USA, E.

## Abstract

Pannexin 1 (Panx1) forms high conductance channels that secrete ATP upon stimulation. The role of Panx1 in mediating constriction in response to direct sympathetic nerve stimulation is not known. Additionally, it is unknown how the expression level of Panx1 in SMCs influences a-adrenergic responses. We hypothesized that the amount of Panx1 in SMCs dictates the levels of sympathetic constriction and blood pressure. To test this hypothesis, we used genetically modified mouse models enabling expression of Panx1 in vascular cells to be varied. Genetic deletion of SMC Panx1 prevented constriction by electric field stimulation of sympathetic nerves. Conversely, over-expression of Panx1 in SMCs using a ROSA26 transgenic model increased sympathetic nerve-mediated constriction. Cx43 hemichannel inhibitors did not alter constriction. Next, we evaluated the effects of altered SMC Panx1 expression on blood pressure. To do this, we created mice combining a global Panx1 deletion, with ROSA26-Panx1 under the control of an inducible SMC specific Cre (Myh11). This resulted in mice that could express only human Panx1, only in SMCs. After tamoxifen, these mice had increased blood pressure that was acutely decreased by the Panx1 inhibitor spironolactone. Control mice genetically devoid of Panx1 did not respond to spironolactone. These data suggest Panx1 in SMCs could regulate the extent of sympathetic nerve constriction and blood pressure. The results also show the feasibility humanized Panx1-mouse models to test pharmacological candidates.

## Introduction

The pannexins are a family of 3 membrane proteins (Panx1, Panx2, and Panx3) that form high conductance channels. In the resistance vasculature, Panx1 is expressed in both endothelial cells (EC) and smooth muscle cells (SMCs).^1^ In SMC, Panx1 channel opening by α_1_-adrenergic stimulation is one of the best described physiological mechanisms to open the channel as demonstrated by electrophysiology, ATP release, and pressure myography of resistance arteries using genetic models.^2-9^ Conditional genetic deletion of Panx1 from SMCs prevents phenylephrine stimulated constriction of resistance arteries. In contrast, genetic deletion of EC Panx1 does not affect systemic hemodynamics, but does regulate cerebral myogenic tone.^2, 10^ As one might expect from these observations, the deletion of Panx1 in SMCs results in a reduction of blood pressure.^2, 3^

Pharmacological approaches to Panx1 inhibition are difficult to interpret and imprecise. For example, probenecid is an accepted Panx1 inhibitor, however it must be used in mM concentrations, is known to block a plethora of other ion channels, and more recently was unable to be resolved in high resolution structures of Panx1.^11^ One of the most specific Panx1 inhibitor with single channel efficacy is spironolactone. However, this drug has been used for decades to block the mineral corticoid receptor chronically. As such, making broad assumptions on Panx1 channels using any one drug can be misleading and difficult to interpret. One method to define a role more precisely for Panx1, and possibly compliment the pharmacology, is the use of genetically modified animals.

While it is known that Panx1 contributes to α_1_-adrenergic stimulated vasoconstriction, its contribution to vasoconstriction in response to direct sympathetic stimulation has not been examined. Additionally, it remains unknown how the expression level of Panx1 in SMCs influences the response to sympathetic stimulation and blood pressure. We hypothesized that the amount of Panx1 in SMCs dictates the levels of sympathetic vasoconstriction and blood pressure. More specifically, that genetic deletion of Panx1 from SMCs impairs sympathetic vasoconstriction and lowers blood pressure, while genetic over-expression of Panx1 in SMCs increases sympathetic vasoconstriction and blood pressure. We tested this hypothesis using a series of mouse models that enabled Panx1 expression to be toggled genetically in SMCs.

## Materials and Methods

### Mice

*Panx1*^fl/fl^ were used as controls and to create inducible smooth muscle knockouts (*Panx1*^fl/fl^ / *Myh11*-CreER^T2+^ as previously described^3^; iSMC Panx1 KO) or inducible endothelial cells knockouts (*Panx1*^fl/fl^ / *Cdh5*-CreER^T2+^ as previously described^12^; iEC Panx1 KO). The ROSA26-Panx1^TG^ mouse with LoxP sites flanking a STOP codon was previously utilized by us to over-express human Panx1.^7^ Like mouse Panx1, human Panx1 is opened by α-adrenergic activation as demonstrated by single-channel recordings and human resistance arteries (e.g., ^2^). These mice were breed with the *Myh11*-CreER^T2+^ to create mice with over-expression of Panx1 in SMC upon tamoxifen induction (ROSA26-*Panx1*^TG^ / *Myh11*-CreER^T2+^; iSMC Panx1 OE). The ROSA26-*Panx1*^TG^ / *Myh11*-CreER^T2+^ mice were then bred with Panx1^−/−^ mice (Panx1^−/−^ mice were originally described^13^; this cross created mice lacking all endogenous Panx1, but inducible expression of Panx1 only in SMC (*Panx1*^−/−^ / ROSA26-*Panx1*^TG^ / *Myh11*-CreER^T2+^; Panx1KO-iSMC OE). In all cases, only male mice were used because the Myh11 Cre transgene, which is specific to smooth muscle and does not target cardiac or skeletal muscle, is found exclusively on the Y-chromosome. Littermates were used in all instances. See **Table1** for complete list of mice used in the experiments described herein, as well as their abbreviations in the figures.

**Table 1:**
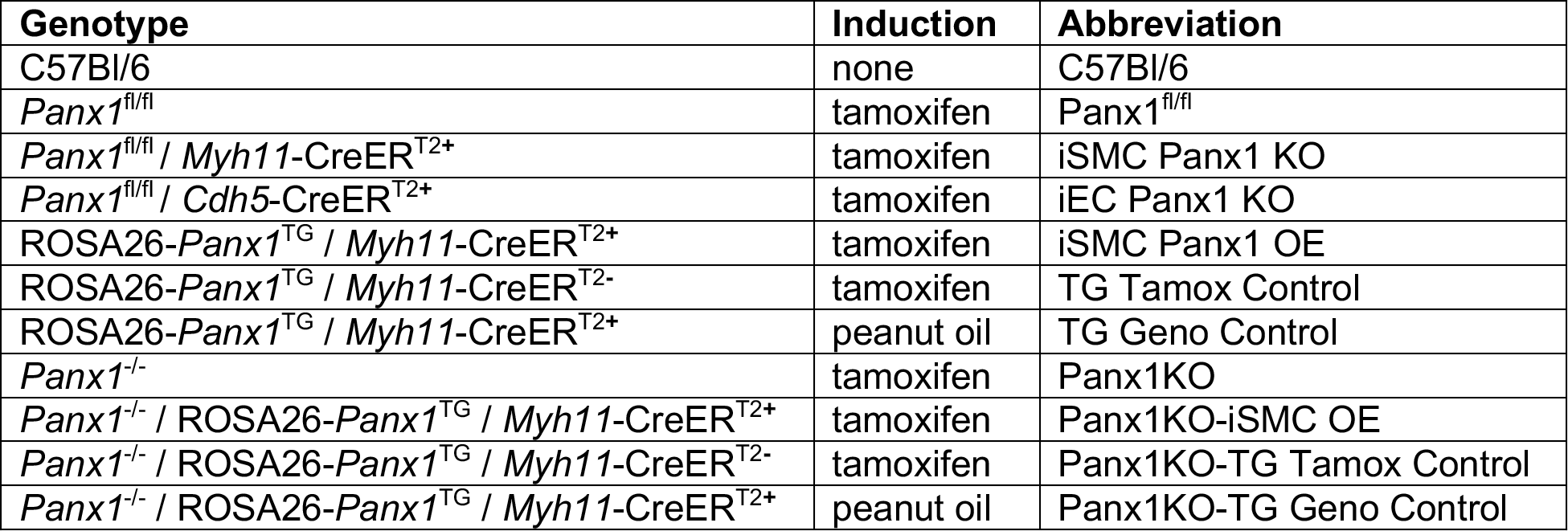
Genotype abbreviations of mice used in this manuscripts’ experiments.

| Genotype | Induction | Abbreviation |
| --- | --- | --- |
| C57Bl/6 | none | C57Bl/6 |
| *Panx1*<sup>fl/fl</sup> | tamoxifen | *Panx1*<sup>fl/fl</sup> |
| *Panx1*<sup>fl/fl</sup> / *Myh11*-CreER<sup>T2+</sup> | tamoxifen | iSMC *Panx1* KO |
| *Panx1*<sup>fl/fl</sup> / *Cdh5*-CreER<sup>T2+</sup> | tamoxifen | iEC *Panx1* KO |
| ROSA26-*Panx1*<sup>TG</sup> / *Myh11*-CreER<sup>T2+</sup> | tamoxifen | iSMC *Panx1* OE |
| ROSA26-*Panx1*<sup>TG</sup> / *Myh11*-CreER<sup>T2-</sup> | tamoxifen | TG Tamox Control |
| ROSA26-*Panx1*<sup>TG</sup> / *Myh11*-CreER<sup>T2+</sup> | peanut oil | TG Geno Control |
| *Panx1*<sup>-/-</sup> | tamoxifen | *Panx1*KO |
| *Panx1*<sup>-/-</sup> / ROSA26-*Panx1*<sup>TG</sup> / *Myh11*-CreER<sup>T2+</sup> | tamoxifen | *Panx1*KO-iSMC OE |
| *Panx1*<sup>-/-</sup> / ROSA26-*Panx1*<sup>TG</sup> / *Myh11*-CreER<sup>T2-</sup> | tamoxifen | *Panx1*KO-TG Tamox Control |
| *Panx1*<sup>-/-</sup> / ROSA26-*Panx1*<sup>TG</sup> / *Myh11*-CreER<sup>T2+</sup> | peanut oil | *Panx1*KO-TG Geno Control |

At 6 weeks of age, all mice were injected daily with 1mg of tamoxifen (mixed in peanut oil as vehicle) for 10 consecutive days (up to 8 weeks in age), and then allowed a two-week tamoxifen washout period so that all experiments were performed on mice at the minimum of 10 weeks of age, up to 15 weeks of age. Mice had a 12h light/dark cycle, were kept at 22-24°C, and had access to water and food *ad libitum*. Mice were used for experiments following the guidelines of the University of Virginia Animal Care and Use Committee.

### Immunocytochemistry

Immunocytochemistry of mesenteric cross-sections from mice were performed as previously described with Panx1 antibody (Cell Signaling 91137).^14^

### Western Blots

Western blots of isolated mesenteric arteries were performed as previously described.^15^ Antibodies used were Panx1 (Cell Signaling 91137) and beta-actin (Cell Signaling 8457). Total protein for blots was provided by Li-Cor Corporation.

### ATP Assays

Both ATP release and intracellular ATP from isolated mesenteric arteries were performed using a ATP Bioluminescence HSII kit (Roche) kit as previously described.^3^

### Electrical Field Stimulation (EFS)

Third order mesenteric arteries were isolated for electrical field stimulation based on previously reported methods.^16^ Sympathetic nerves on third order mesenteric arteries were stimulated by two platinum electrodes in a bath chamber (STIM-1501; Living Systems Instrumentation). To measure sympathetic vasoconstriction electrical pulses (70 V, 2 ms) were delivered at 4, 8, 12, and 16 Hz. Pulses were performed every 5 minutes.

### Urinary norepinephrine (NE)

Urine was collected from mice in metabolic cages after a two-week adjustment period. A norepinephrine ELISA (KA1877, Novus) was used to quantify amounts.

### Radiotelemetry

Radiotelemetry procedures were performed as previously described.^16^

### Statistics

Shapiro-Wilks normality of data was established in all cases. One- or two-way ANOVA’s as well as two tailed t-tests were performed where indicated using GraphPad Prism version 9.3.1. Data is represented as the mean ± SEM with a p-value of <0.05 used as the threshold for statistical significance. For analyses requiring two-way ANOVA, Holm-Sidak or Sidak post hoc analysis were used to compare groups.

## Results

We have previously shown that phenylephrine (PE)-stimulated constriction of arteries is blunted by pharmacological and genetic deletion of Panx1 in SMC^3, 5^. We therefore hypothesized sympathetic nerve-induced constriction may be regulated by the amount of Panx1 that resides in SMC. To test this, we created a mouse with inducible over-expression of Panx1 using a ROSA26 insert^7^ that was crossed with *Myh11*CreER^T2+^ mice (iSMC Panx1 OE mice). In response to tamoxifen treatment, there was a significant increase in SMC Panx1 expressed by iSMC Panx1 OE mice (**Figure 1A-B; Supplemental Figure 1**).

**Figure 1:**
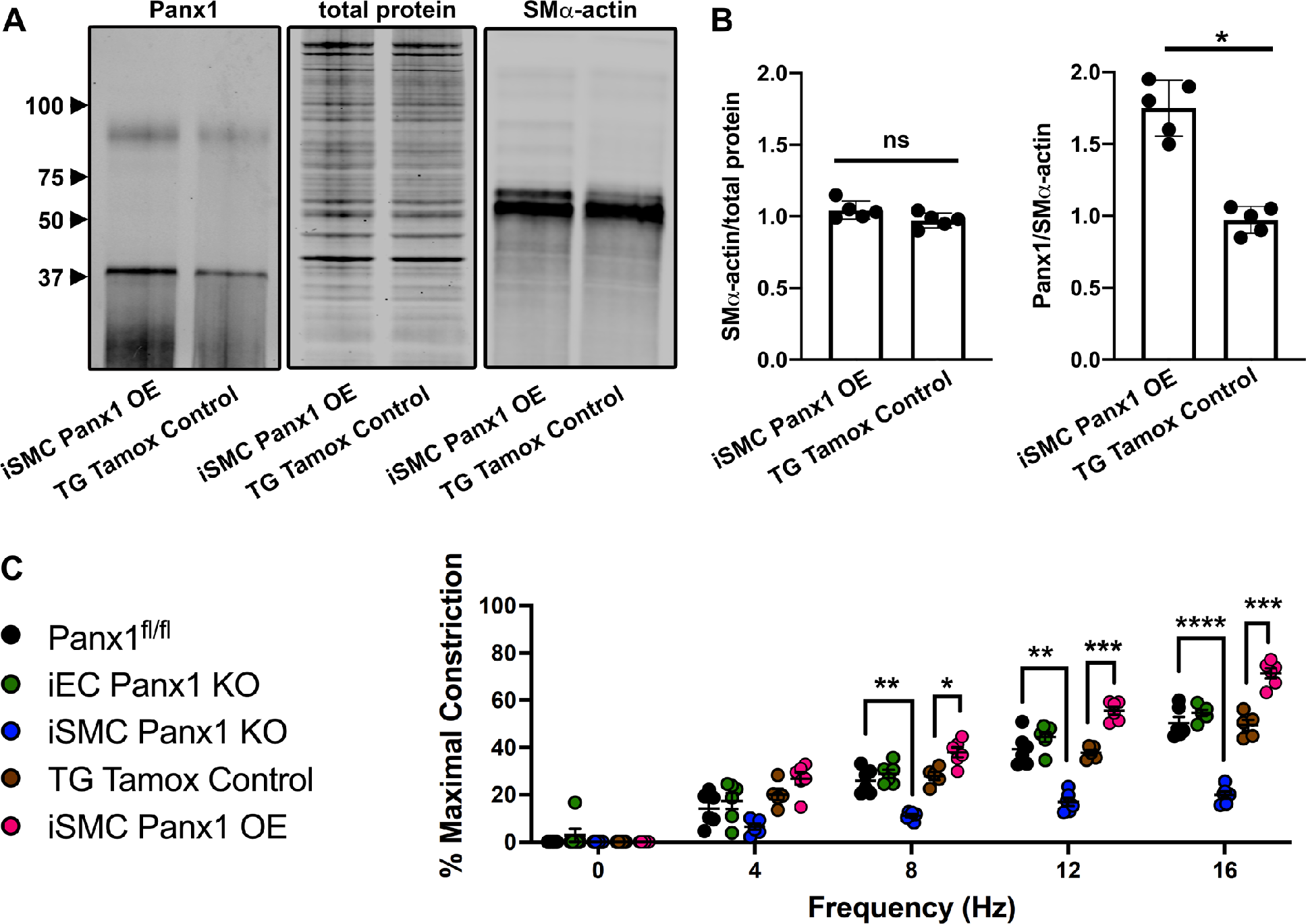
Over-expression of Panx1 in smooth muscle cells increases vasoconstriction after sympathetic nerve stimulation. (**A-B**) Western blot and quantification of Panx1 abundance in mesenteric arteries of iSMC Panx1 OE and TG Tamox Control mice * p<0.05. (**C**) EFS-induced constriction of third order mesenteric arteries from Panx1^fl/fl^, iEC Panx1 KO, and iSMC Panx1 KO mice. **p<0.01, ***p<0.001, **** p<0.0001 Panx1^fl/fl^ vs iSMC Panx1 KO. Data were analyzed by repeated measures two-way ANOVA and Sidak post hoc test.

To test whether vasoconstriction induced by sympathetic nerve stimulation correlated with differences in SMC Panx1 expression, we used electric field stimulation (EFS) of third order mesenteric arteries from iSMC Panx1 OE mice. In comparison with control mice, there was a significant increase in vasoconstriction (**Figure 1C**). In contrast, inducible mice lacking SMC Panx1 (iSMC Panx1 KO) had a significantly decreased response to EFS. Confirming a lack of EC Panx1 contribution to systemic sympathetic-driven vasoconstriction, inducible mice lacking EC Panx1 expression (iEC Panx1 KO) had a response to EFS that was comparable to control mice expressing normal levels of EC Panx1 (**Figure 1C**). There were no differences in EFS-induced constriction in any of the control groups examined (Panx1^fl/fl^, TG Tamox Control, TG Geno Control) (**Supplemental Figure 2**).

To confirm EFS induced constriction was mediated by adrenergic nerve stimulation, we blocked norepinephrine release with guanethidine (10 μM) and blocked α-adrenergic receptors with phentolamine (1 μM). We found both guanethidine and phentolamine blunted EFS-induced constriction in all genotypes except for iSMC Panx1 OE (**Supplement Figure 3-4**) demonstrating the Panx1 activation by SMCs was downstream from α-adrenergic stimulation.

ATP release is a hallmark of Panx1 channel opening. We therefore investigated ATP release in mice with variable genetic expression of Panx1 in SMC by stimulating isolated third order mesenteric arteries with 10 μM PE. We found iSMC Panx1 KO mice to have significantly blunted ATP release and iSMC OE mice to have significantly increased ATP release compared to their respective controls (**Figure 2A**). These results correlate levels of SMC Panx1 expression with levels of ATP release. Mice lacking Panx1 in EC had no change in ATP release in response PE. Importantly, ATP vessel content was comparable in all the mouse models examined, indicating that strain specific differences in ATP release were due to differences in secretion (**Supplemental Figure 5**). Because PE stimulates ATP release from third order mesenteric arteries in a Panx1-dependent manner, we hypothesized EFS induces constriction in an ATP-dependent manner. To test this, we hydrolyzed extracellular ATP by adding 50 U/mL apyrase to vessels undergoing EFS. Apyrase blunted EFS-induced constriction in Panx1^fl/fl^, iEC Panx1 KO, and iSMC Panx1 OE mice, but not iSMC Panx1 KO mice. (**Figure 2B-E**).

**Figure 2:**
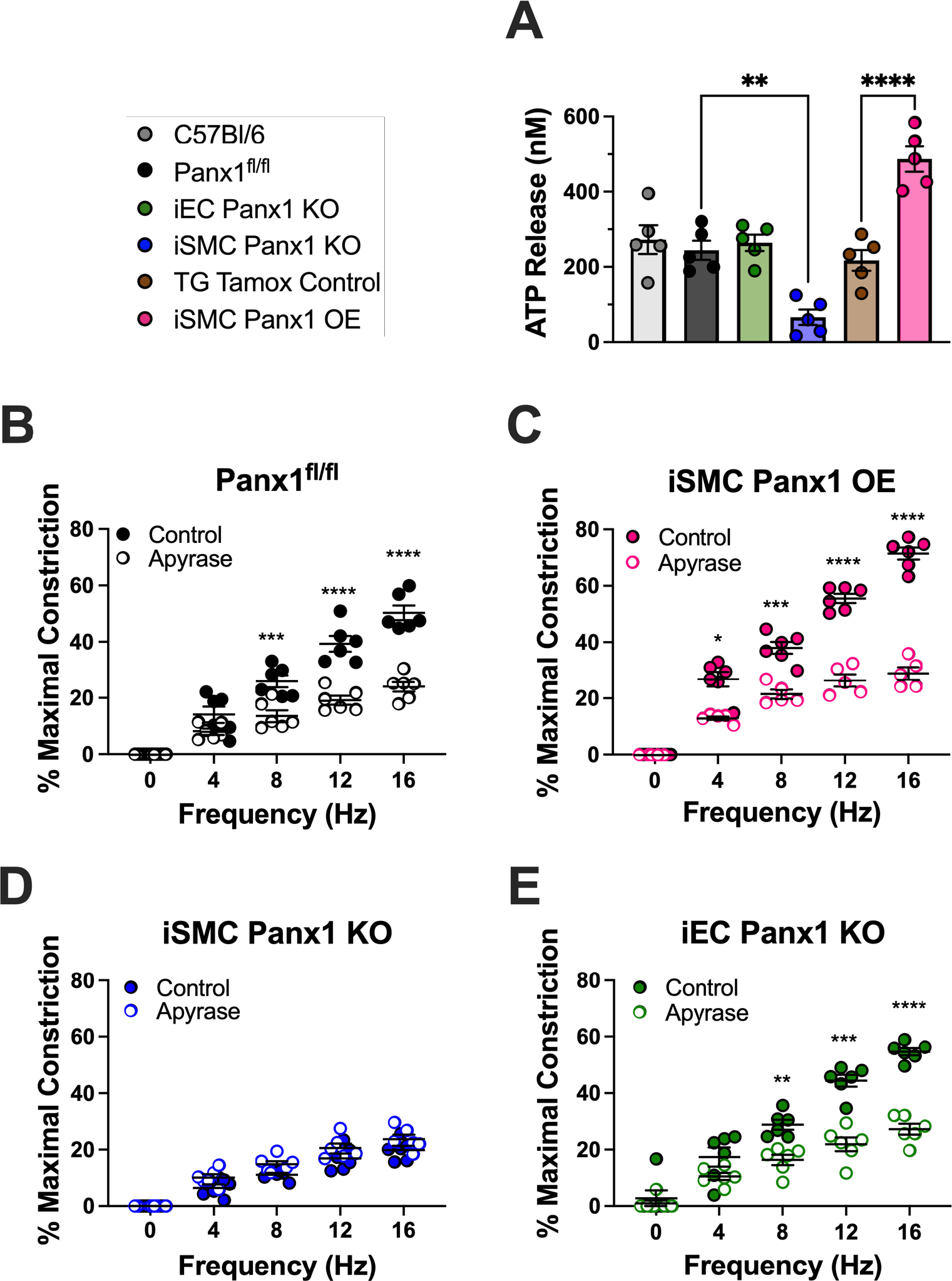
Increased ATP release from mice over-expressing Panx1 in smooth muscle cells. (**A**) ATP release from third order mesenteric arteries after stimulation with 10 μM phenylephrine. Data were analyzed by one-way ANOVA and Holm-Sidak post hoc test. ** p<0.01; *** p<0.0001 (**B-E**) EFS-induced constriction of third order mesenteric arteries with or without apyrase (50 U/mL). Data were analyzed by repeated measures two-way ANOVA and Sidak post hoc test. *p<0.05, **p<0.01, ***p<0.001, **** p<0.0001

We further investigated the role of Panx1 in EFS induced constriction using pharmacological agents (**Figure 3**) and peptides (**Figure 4**) known to inhibit Panx1. We found that 2 mM probenecid^17^ and 20 μM spironolactone^2^ both blunted EFS-induced constriction of vessels from Panx1^fl/fl^, iEC Panx1 KO, and iSMC Panx1 OE mice, but not iSMC Panx1 KO mice (**Figure 3** and **Supplemental Figure 6**). We have previously shown that the internal loop of Panx1 is important for Panx1 activation and localization and that a peptide recognizing this region (PxIL2P) is efficacious in blocking the channel ^3, 14, 15, 18^. PxIL2P (20 μM) blunted constriction in Panx1^fl/fl^, iEC Panx1 KO, and iSMC Panx1 OE mice, but not iSMC Panx1 KO mice, comparable with results obtained using pharmacologic inhibitors of the channel (**Figure 4A-C and Supplemental Figure 6**). As a control to rule out ATP secretion by connexins, we found that the Cx43 peptide inhibitor Gap19 did not affect EFS-induced constriction (**Figure 4D-F and Supplemental Figure 6**).

**Figure 3:**
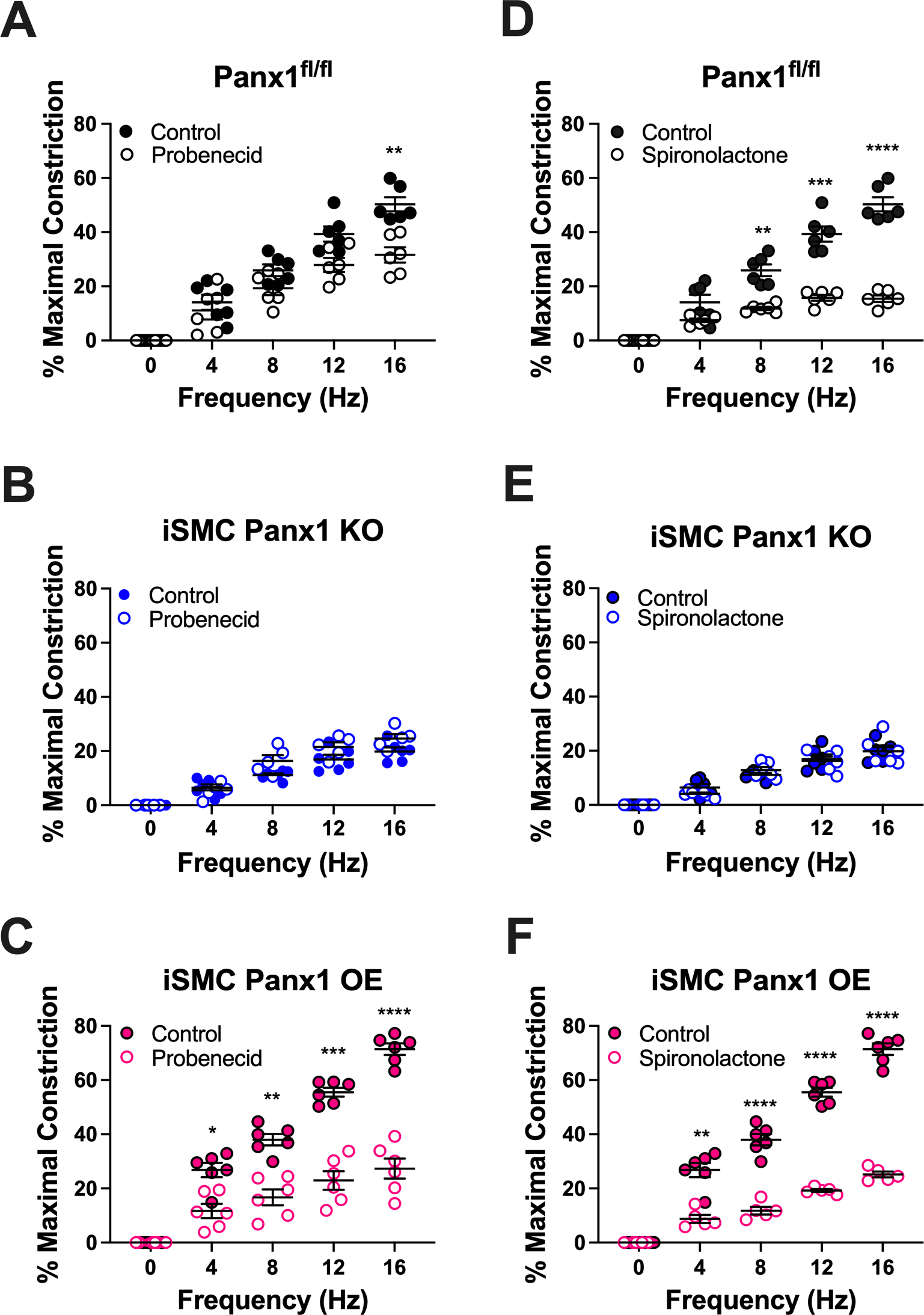
Pharmacological inhibition of Panx1 inhibits sympathetic nerve induced vasoconstriction. (**A-C**) EFS-induced vasoconstriction with or without probenecid (2 mM) (**D-F**) EFS-induced vasoconstriction with or without spironolactone (20 μM). Data were analyzed by repeated measures two-way ANOVA and Sidak post hoc test. *p<0.05, **p<0.01, ***p<0.001, **** p<0.0001

**Figure 4:**
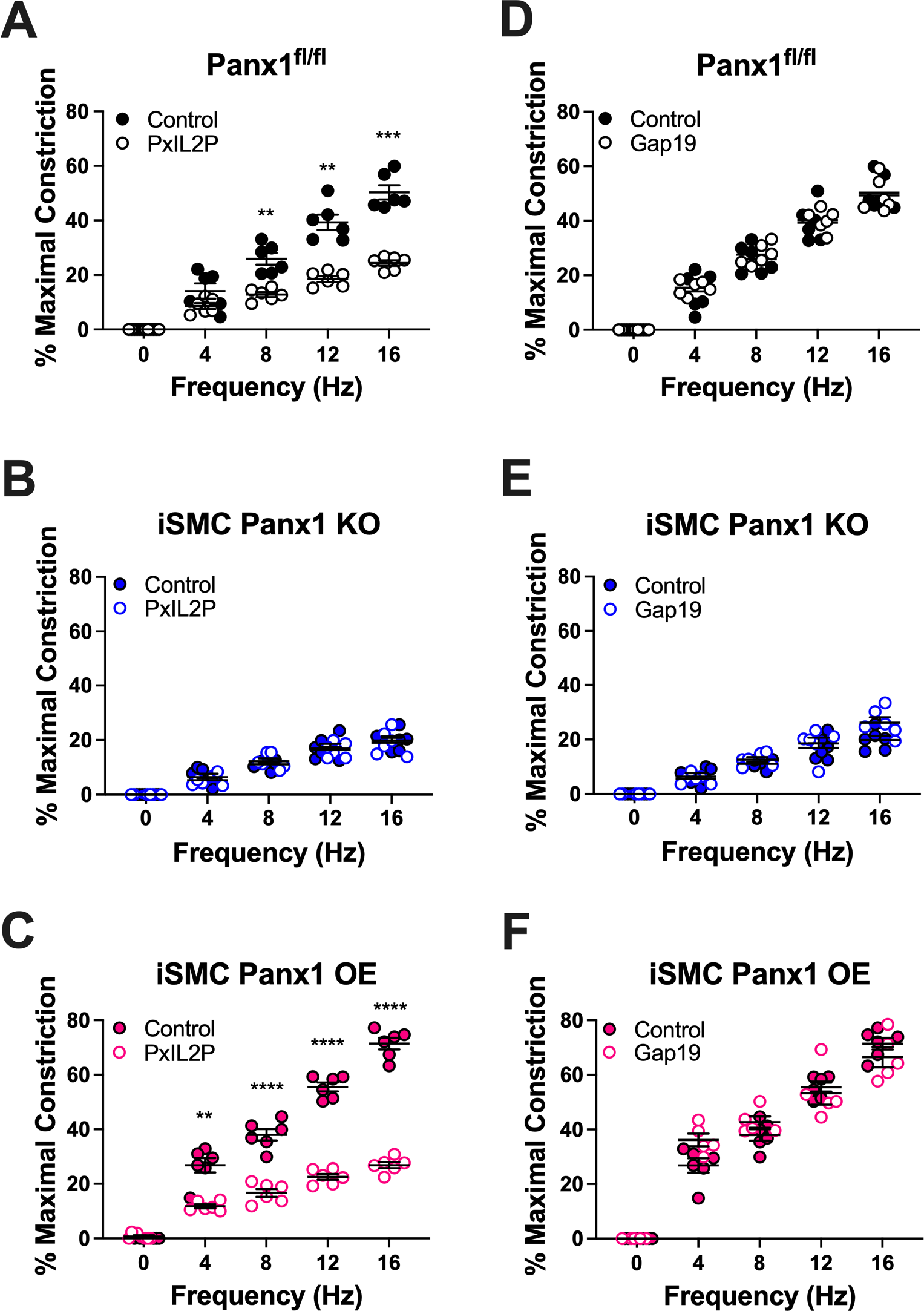
Peptide inhibition of Panx1, but not Cx43, inhibits sympathetic nerve vasoconstriction. (**A-C**) EFS-induced constriction with or without the Panx1 inhibitor, PxIL2P (20 μM) or (**D-F**) EFS-induced constriction with or without the Cx43 inhibitor, Gap19 (20 μM). Data were analyzed by repeated measures two-way ANOVA and Sidak post hoc test. **p<0.01, ***p<0.001, **** p<0.0001

Genetic Panx1 deletion has been used to demonstrate an important role in blood pressure regulation by renin-secreting cells^18^ (where genetic deletion causes hypertension) and by SMC^3^ (where genetic deletion causes hypotension). Because iSMC Panx1 OE mice displayed increased EFS induced vasoconstriction we hypothesized these mice would have elevated blood pressure. Blood pressure was measured prior to injection of tamoxifen or peanut oil and again after treatment as indicated in **Table 1**. Upon induction with tamoxifen (or peanut oil), only iSMC Panx1 OE mice showed a significant increase in blood pressure compared to the pre-induction baseline (**Figure 5A**). None of the other mouse models had altered blood pressure. By contrast, acute inhibition of Panx1 by intraperitoneal injection of 40 mg/kg spironolactone resulted in a drop in blood pressure in iSMC Panx1 OE mice and transgenic controls which express endogenous SMC Panx1 (**Figure 5A**). Note that all the mice had any similar levels of urinary norepinephrine (**Supplemental Figure 7**).

**Figure 5:**
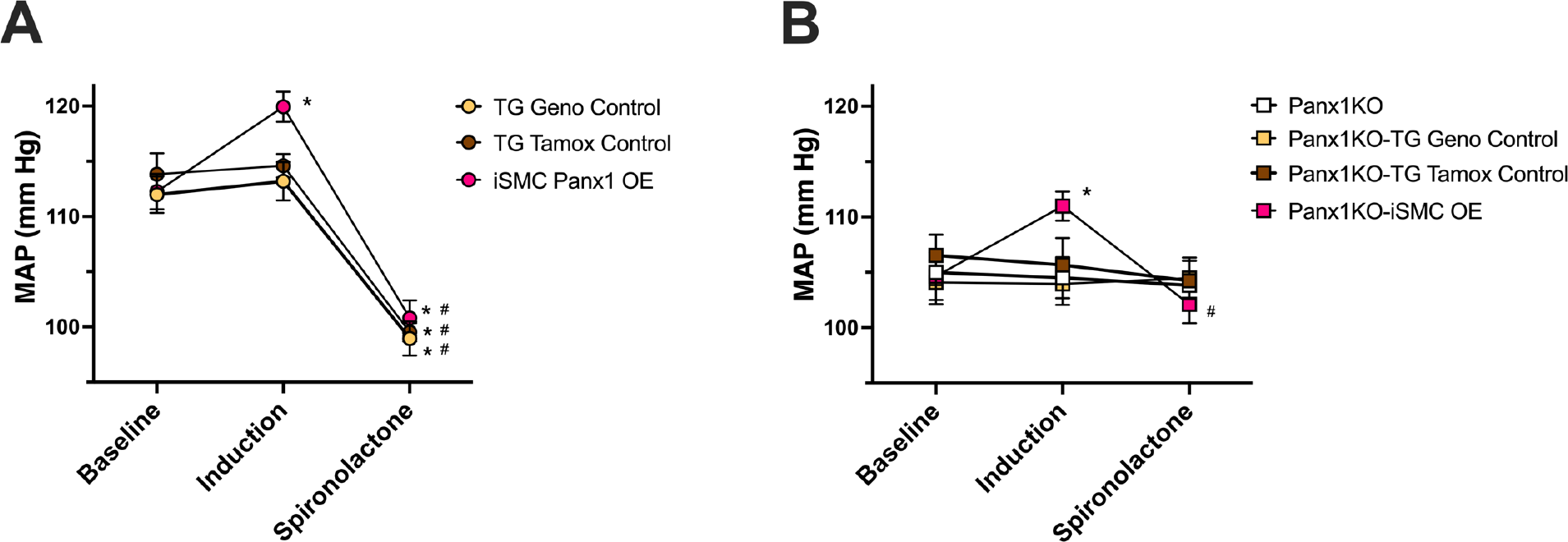
Overexpression of human Panx1 in SMC alone increases blood pressure and is rescued by acute injection of spironolactone. Blood pressure was recorded at baseline, after treatment with tamoxifen or vehicle (induction), or after acute treatment with 40 mg/kg spironolactone via intraperitoneal injection. (**A**) Mice with baseline level of Panx1 (yellow, TG Geno Control and TG Tamox Control, brown) and after induction, had over-expression of Panx1 (red, iSMC Panx1 OE). (**B**) Mice were baseline lacking Panx1 globally (white, Panx1KO; yellow Panx1KO-TG Geno Control; brown, Panx1KO-TG Tamox Control) and after induction, only had Panx1 in SMC (red, Panx1KO-iSMC OE). Data were analyzed by two-way ANOVA and Holm-Sidak post hoc test. * p<0.05 vs respective baseline; # p<0.05 vs respective induction.

We further investigated a role for Panx1 in regulating blood pressure by using global Panx1 knockout mice (*Panx1*^−/−^). To test only the effect of Panx1 in SMC and no other cell types, we crossed iSMC Panx1 OE mice onto a Panx1^−/−^ background, enabling us to evaluate blood pressure in mice exclusively expressing hPanx1 in SMCs (Panx1KO-iSMC OE; genotypes shown in **Table 1)**. Panx1KO-iSMC OE mice had increased blood pressure after induction with tamoxifen. No difference in blood pressure was observed in control mice which received either tamoxifen (Panx1KO-TG Tamox Control) or peanut oil (Panx1KO-TG Geno Control). Moreover, acute inhibition of Panx1 with spironolactone was only able to reduce blood pressure in Panx1KO-iSMC OE mice (**Figure 5B**), further supporting a role for SMC Panx1 in sympathetic nerve-mediated changes in blood pressure.

## Discussion

We have previously demonstrated that SMC Panx1 has the capacity to regulate blood pressure.^2, 3, 18^ Here we build on this work using mouse models that enable Panx1 expression by SMCs to be further regulated, essentially enabling a dose response of Panx1 to be measured. Our findings demonstrate that sympathetic vasoconstriction and blood pressure are dependent, at least to some extent, on the amount of Panx1 expressed by SMCs.

We observed a blunted response to EFS-mediated constriction in iSMC Panx1 KO mice, consistent with previous studies demonstrating decreased constriction in response to α_1_-adrenergic stimulation.^3^ We additionally show iSMC Panx1 OE mice have increased EFS-mediated constriction. This occurs through sympathetic nerve stimulation as demonstrated by the ability of guanethidine and phentolamine to inhibit EFS-mediated constriction. iEC Panx1 KO mice responded similarly to control mice suggesting SMC Panx1, and not EC Panx1, regulates sympathetic vasoconstriction. Although mice lacking EC Panx1 have alterations in cerebral myogenic tone, no systemic hemodynamic phenotype has been observed.^2, 10^ This is consistent with a central role for SMC Panx1 in regulating systemic blood pressure.

Based on the experiments reported here and our previous work, we propose the increased vasoconstriction in response to EFS is due, at least in part, to the release of ATP from the increased number of Panx1 channels in the SMC. ATP release from Panx1 channels has been extensively demonstrated (e.g.^19-23^), and α_1_-adrenergic stimulation is one of the most well characterized mechanisms to activate and open Panx1 channels (e.g.^2-9^). This model is strongly supported by our findings that iSMC Panx1 OE mice have greater ATP release after PE stimulation and the hydrolysis of ATP by apyrase blunts EFS constriction. However, we cannot completely exclude two other possible Panx1-dependent mechanisms to regulate sympathetic vasoconstriction and blood pressure. The first possibility is that changes in Panx1 by SMCs may alter sympathetic nerve activity through a yet to be determined mechanism—possibly SMC intracellular calcium concentrations in SMC, or vesicular nucleotide transporter (VNUT) ATP activity in sympathetic nerves. While we did not directly investigate these possibilities, we found changes in urinary norepinephrine were unaltered across all the genotypes examined, which suggest at least sympathetic nerve activity is unaltered by Panx1 in SMCs. The second possibility is that the vasoconstriction may be due to the capacity of Panx1 channels to facilitate anion transport across the plasma membrane, e.g. Cl^−^.^24^ Although our observation that apyrase blunts EFS constriction suggests a role for extracellular ATP in vasoconstriction, we cannot completely rule out a possible additional contribution of anions such as Cl^−^, since changes in Cl^−^ are a well-known mechanism to induce SMC vasoconstriction^25^ and are worthy of consideration and future study for Panx1 channel activity.

To the best of our knowledge, this is the first time a mouse with only the Panx1 sequence from human was utilized (Panx1KO-iSMC OE). Like many mouse models where the endogenous murine proteins have been replaced by the human protein equivalent (“humanized mouse”), there is strong utility in this type of model system (e.g., ^26^). In our system we tested whether human Panx1 in smooth muscle alone could alter blood pressure or respond to the Panx1 spironolactone in an intact system. In both cases the mice responded when Panx1 was only present in SMC, and nowhere else in the body. These experiments lay the groundwork for many other studies to not only test the specific role of Panx1 when expressed in only one cell type, but the physiological effects of potential pharmacological interventions using the human form of Panx1.

We demonstrated a role in SMC Panx1 control of blood pressure. We have previously observed a decrease in blood pressure in iSMC Panx1 KO mice as well as accompanying acute inhibition of Panx1 with spironolactone.^2, 3^ Here we found that overexpression of SMC Panx1 increased blood pressure regardless of levels of systemic Panx1 expression. Additionally, acute inhibition of Panx1 with spironolactone reduced blood pressure only in mice expressing Panx1 in SMCs, further underscoring the importance of Panx1 expressed by these cells in regulating blood pressure.

Spironolactone is a well-recognized antihypertensive agent shown to target mineralocorticoid receptors. We recently demonstrated spironolactone also has the ability to inhibit Panx1 channels.^2^ While chronic blood pressure reduction with spironolactone is primarily mediated through mineralocorticoid antagonism, our data using multiple mouse models to specifically regulate Panx1 expression by SMCs further supports the conclusion that Panx1 inhibition is a critical element of the acute reduction in blood pressure by spironolactone. In addition to adding support for the use of spironolactone to control blood pressure, our data underscore the importance of identifying and considering other agents that target Panx1 in therapeutic approaches to regulate blood pressure.

We complemented our mouse models using pharmacological inhibitors of Panx1 including probenecid, spironolactone, and PxIL2P. Acute inhibition of Panx1 ex vivo blunted EFS-mediated constriction in all groups except iSMC Panx1 KO mice, further supporting the conclusion that SMC Panx1 plays an active role in sympathetic vasoconstriction. However, it is not clear whether Panx1 itself is a good pharmacological target due to its ubiquitous expression. For example, deletion of Panx1 in renin-secreting cells induces an increase in blood pressure whereas deletion of Panx1 in SMCs induces a reduction in blood pressure. Thus, any pharmacological intervention may have a net-zero effect on blood pressure. For example, recent studies have indicated that probenecid has no effect on blood pressure^27^, albeit at significantly lower concentrations than those required to inhibit Panx1 channels^28^. However, using genetic models we have been able to parse out individual contributions of cell types by selectively deleting or increasing the amount of Panx1 expressed by SMCs and have shown Panx1 can serve as a rheostat to regulate blood pressure. The work from these findings supports targeting cell type-specific Panx1 expression and ATP release as a novel approach to the control of blood pressure.

Together the data presented here suggest the amount of Panx1 in SMC dictates the magnitude of constriction in response to sympathetic stimulation and consequently regulates blood pressure levels. This raises the possibility that Panx1 expression levels could represent a risk factor for hypertension. If this is the case, a screen for differential Panx1 expression could have diagnostic value. In addition, strategies that target levels of SMC Panx1 expression may have the capacity to complement approaches that modulate Panx1 channel activity to control blood pressure.

## Acknowledgements

We thank the pannexin interest group at the University of Virginia for critical feedback in development and implementation of the mice.

## Sources of Funding

National Institutes of Health (NIH) grants T32 007284 (LSD), HL 120840 (KR, UL, and BEI), and HL 137112 (MK and BEI).

## Disclosures

None.

## Abbreviations

Panx1: pannexin 1
SMC: Smooth muscle cell
PE: phenylephrine
EC: endothelial cell
EFS: electrical field stimulation

## Notes

### Competing Interest Statement

The authors have declared no competing interest.

## References

1. Lohman AW, Billaud M, Straub AC, Johnstone SR, Best AK, Lee M, Barr K, Penuela S, Laird DW and Isakson BE. Expression of pannexin isoforms in the systemic murine arterial network. J Vasc Res. 2012;49:405–16.

2. Good ME, Chiu YH, Poon IKH, Medina CB, Butcher JT, Mendu SK, DeLalio LJ, Lohman AW, Leitinger N, Barrett E, Lorenz UM, Desai BN, Jaffe IZ, Bayliss DA, Isakson BE and Ravichandran KS. Pannexin 1 Channels as an Unexpected New Target of the Anti-Hypertensive Drug Spironolactone. Circ Res. 2018;122:606–615.

3. Billaud M, Chiu YH, Lohman AW, Parpaite T, Butcher JT, Mutchler SM, DeLalio LJ, Artamonov MV, Sandilos JK, Best AK, Somlyo AV, Thompson RJ, Le TH, Ravichandran KS, Bayliss DA and Isakson BE. A molecular signature in the pannexin1 intracellular loop confers channel activation by the alpha1 adrenoreceptor in smooth muscle cells. Sci Signal. 2015;8:ra17.

4. DeLalio LJ, Keller AS, Chen J, Boyce AKJ, Artamonov MV, Askew-Page HR, Keller TCSt, Johnstone SR, Weaver RB, Good ME, Murphy SA, Best AK, Mintz EL, Penuela S, Greenwood IA, Machado RF, Somlyo AV, Swayne LA, Minshall RD and Isakson BE. Interaction Between Pannexin 1 and Caveolin-1 in Smooth Muscle Can Regulate Blood Pressure. Arterioscler Thromb Vasc Biol. 2018;38:2065–2078.

5. Billaud M, Lohman AW, Straub AC, Looft-Wilson R, Johnstone SR, Araj CA, Best AK, Chekeni FB, Ravichandran KS, Penuela S, Laird DW and Isakson BE. Pannexin1 regulates alpha1-adrenergic receptor- mediated vasoconstriction. Circulation Research. 2011;109:80–5.

6. Chiu YH, Medina CB, Doyle CA, Zhou M, Narahari AK, Sandilos JK, Gonye EC, Gao HY, Guo SY, Parlak M, Lorenz UM, Conrads TP, Desai BN, Ravichandran KS and Bayliss DA. Deacetylation as a receptor-regulated direct activation switch for pannexin channels. Nat Commun. 2021;12:4482.

7. Medina CB, Chiu YH, Stremska ME, Lucas CD, Poon I, Tung KS, Elliott MR, Desai B, Lorenz UM, Bayliss DA and Ravichandran KS. Pannexin 1 channels facilitate communication between T cells to restrict the severity of airway inflammation. Immunity. 2021;54:1715–1727 e7.

8. Adamson SE, Meher AK, Chiu YH, Sandilos JK, Oberholtzer NP, Walker NN, Hargett SR, Seaman SA, Peirce-Cottler SM, Isakson BE, McNamara CA, Keller SR, Harris TE, Bayliss DA and Leitinger N. Pannexin 1 is required for full activation of insulin-stimulated glucose uptake in adipocytes. Mol Metab. 2015;4:610–8.

9. Tozzi M, Hansen JB and Novak I. Pannexin-1 mediated ATP release in adipocytes is sensitive to glucose and insulin and modulates lipolysis and macrophage migration. Acta physiologica. 2020;228:e13360.

10. Good ME, Eucker SA, Li J, Bacon HM, Lang SM, Butcher JT, Johnson TJ, Gaykema RP, Patel MK, Zuo Z and Isakson BE. Endothelial cell Pannexin1 modulates severity of ischemic stroke by regulating cerebral inflammation and myogenic tone. JCI Insight. 2018;3.

11. Kuzuya M, Hirano H, Hayashida K, Watanabe M, Kobayashi K, Terada T, Mahmood MI, Tama F, Tani K, Fujiyoshi Y and Oshima A. Structures of human pannexin-1 in nanodiscs reveal gating mediated by dynamic movement of the N terminus and phospholipids. Sci Signal. 2022;15:eabg6941.

12. Lohman AW, Leskov IL, Butcher JT, Johnstone SR, Stokes TA, Begandt D, DeLalio LJ, Best AK, Penuela S, Leitinger N, Ravichandran KS, Stokes KY and Isakson BE. Pannexin 1 channels regulate leukocyte emigration through the venous endothelium during acute inflammation. Nat Commun. 2015;6:7965.

13. Poon IK, Chiu YH, Armstrong AJ, Kinchen JM, Juncadella IJ, Bayliss DA and Ravichandran KS. Unexpected link between an antibiotic, pannexin channels and apoptosis. Nature. 2014;507:329–34.

14. Yang Y, Delalio LJ, Best AK, Macal E, Milstein J, Donnelly I, Miller AM, McBride M, Shu X, Koval M, Isakson BE and Johnstone SR. Endothelial Pannexin 1 Channels Control Inflammation by Regulating Intracellular Calcium. J Immunol. 2020.

15. DeLalio LJ, Billaud M, Ruddiman CA, Johnstone SR, Butcher JT, Wolpe AG, Jin X, Keller TCSt, Keller AS, Riviere T, Good ME, Best AK, Lohman AW, Swayne LA, Penuela S, Thompson RJ, Lampe PD, Yeager M and Isakson BE. Constitutive SRC-mediated phosphorylation of pannexin 1 at tyrosine 198 occurs at the plasma membrane. J Biol Chem. 2019;294:6940–6956.

16. Kruger N, Biwer LA, Good ME, Ruddiman CA, Wolpe AG, DeLalio LJ, Murphy S, Macal EH, Jr., Ragolia L, Serbulea V, Best AK, Leitinger N, Harris TE, Sonkusare SK, Godecke A and Isakson BE. Loss of Endothelial FTO Antagonizes Obesity-Induced Metabolic and Vascular Dysfunction. Circ Res. 2020;126:232–242.

17. Sandilos JK, Chiu Y-hH, Chekeni FB, Armstrong AJ, Walk SF, Ravichandran KS and Bayliss DA. Pannexin 1, an ATP release channel, is activated by caspase cleavage of its pore-associated C terminal autoinhibitory region. J Biol Chem. 2012;In Press.

18. DeLalio LJ, Masati E, Mendu S, Ruddiman CA, Yang Y, Johnstone SR, Milstein JA, Keller TCSt, Weaver RB, Guagliardo NA, Best AK, Ravichandran KS, Bayliss DA, Sequeira-Lopez MLS, Sonkusare SN, Shu XH, Desai B, Barrett PQ, Le TH, Gomez RA and Isakson BE. Pannexin 1 Channels in Renin-Expressing Cells Influence Renin Secretion and Blood Pressure Homeostasis. Kidney Int. 2020.

19. Bao L, Locovei S and Dahl G. Pannexin membrane channels are mechanosensitive conduits for ATP. FEBS Lett. 2004;572:65–8.

20. Chekeni FB, Elliott MR, Sandilos JK, Walk SF, Kinchen JM, Lazarowski ER, Armstrong AJ, Penuela S, Laird DW, Salvesen GS, Isakson BE, Bayliss DA and Ravichandran KS. Pannexin 1 channels mediate ‘find-me’ signal release and membrane permeability during apoptosis. Nature. 2010;467:863–7.

21. Billaud M, Lohman AW, Straub AC, Looft-Wilson R, Johnstone SR, Araj CA, Best AK, Chekeni FB, Ravichandran KS, Penuela S, Laird DW and Isakson BE. Pannexin1 regulates alpha1-adrenergic receptor- mediated vasoconstriction. Circ Res. 2011;109:80–5.

22. Gulbransen BD, Bashashati M, Hirota SA, Gui X, Roberts JA, MacDonald JA, Muruve DA, McKay DM, Beck PL, Mawe GM, Thompson RJ and Sharkey KA. Activation of neuronal P2X7 receptor-pannexin-1 mediates death of enteric neurons during colitis. Nature medicine. 2012;18:600–4.

23. Weilinger NL, Tang PL and Thompson RJ. Anoxia-induced NMDA receptor activation opens pannexin channels via Src family kinases. J Neurosci. 2012;32:12579–88.

24. Ruan Z, Orozco IJ, Du J and Lu W. Structures of human pannexin 1 reveal ion pathways and mechanism of gating. Nature. 2020;584:646–651.

25. Wray S, Prendergast C and Arrowsmith S. Calcium-Activated Chloride Channels in Myometrial and Vascular Smooth Muscle. Front Physiol. 2021;12:751008.

26. Walsh NC, Kenney LL, Jangalwe S, Aryee KE, Greiner DL, Brehm MA and Shultz LD. Humanized Mouse Models of Clinical Disease. Annu Rev Pathol. 2017;12:187–215.

27. Gliemann L, Tamariz-Ellemann A, Collin Hansen C, Svare Ehlers T, Moller S and Hellsten Y. Is the Pannexin-1 Channel a Mechanism Underlying Hypertension in Humans? a Translational Study of Human Hypertension. Hypertension. 2022;79:1132–1143.

28. Silverman W, Locovei S and Dahl G. Probenecid, a gout remedy, inhibits pannexin 1 channels. Am J Physiol Cell Physiol. 2008;295:C761–7.

