## Supplemental Figures for "Amount of Pannexin 1 in smooth muscle cells regulates sympathetic nerve induced vasoconstriction"


Supplemental Figures: 7

**Supplemental Figures:**

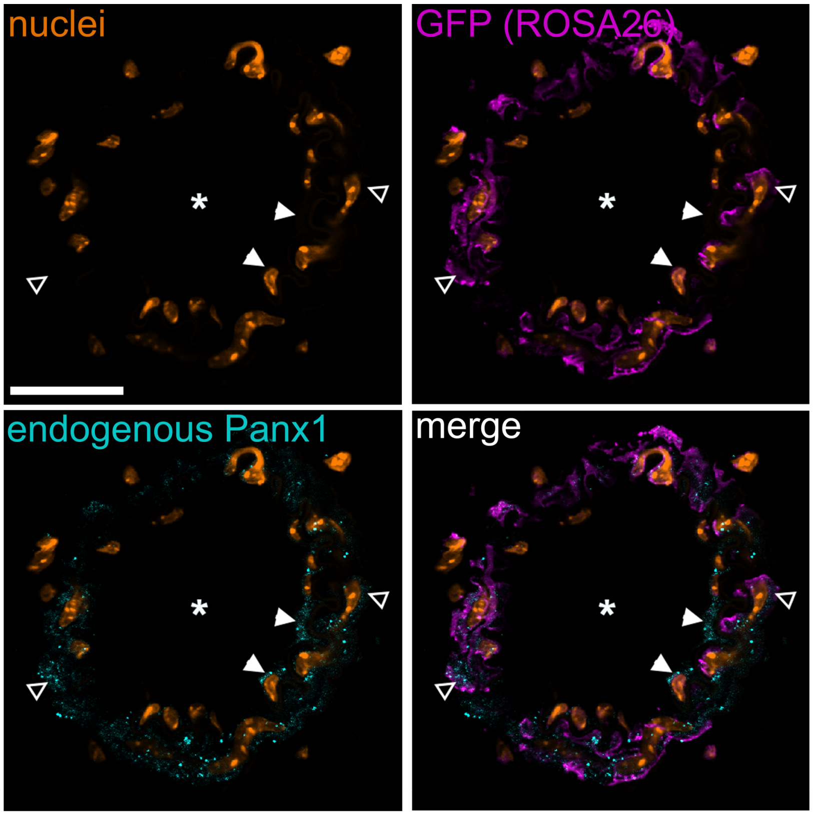


**Supplemental Figure 1: Endogenous and over-expressed Panx1 in iSMC Panx1 OE mesenteric arteries.** In each images, * is lumen of third order mesenteric artery; solid arrowheads point to endothelium, and open arrow heads point to smooth muscle. Nuclei (stained with DAPI) is pseudocolored in orange; GFP from the ROSA26 reporter indicating Panx1 expression is pseudocolored in magenta; endogenous Panx1 is in cyan. Scale bar is 50 μm for each image.

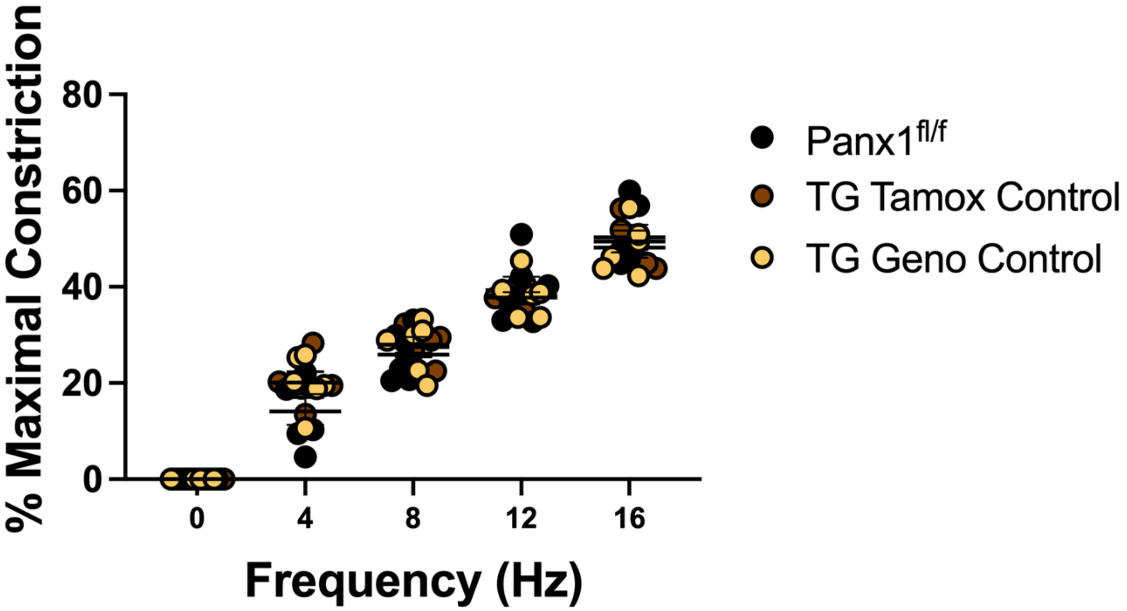


**Supplemental Figure 2: Sympathetic vasoconstriction is similar across control groups.** EFS-induced constriction in control mice. Data was analyzed by two-way ANOVA and Sidak post hoc test.

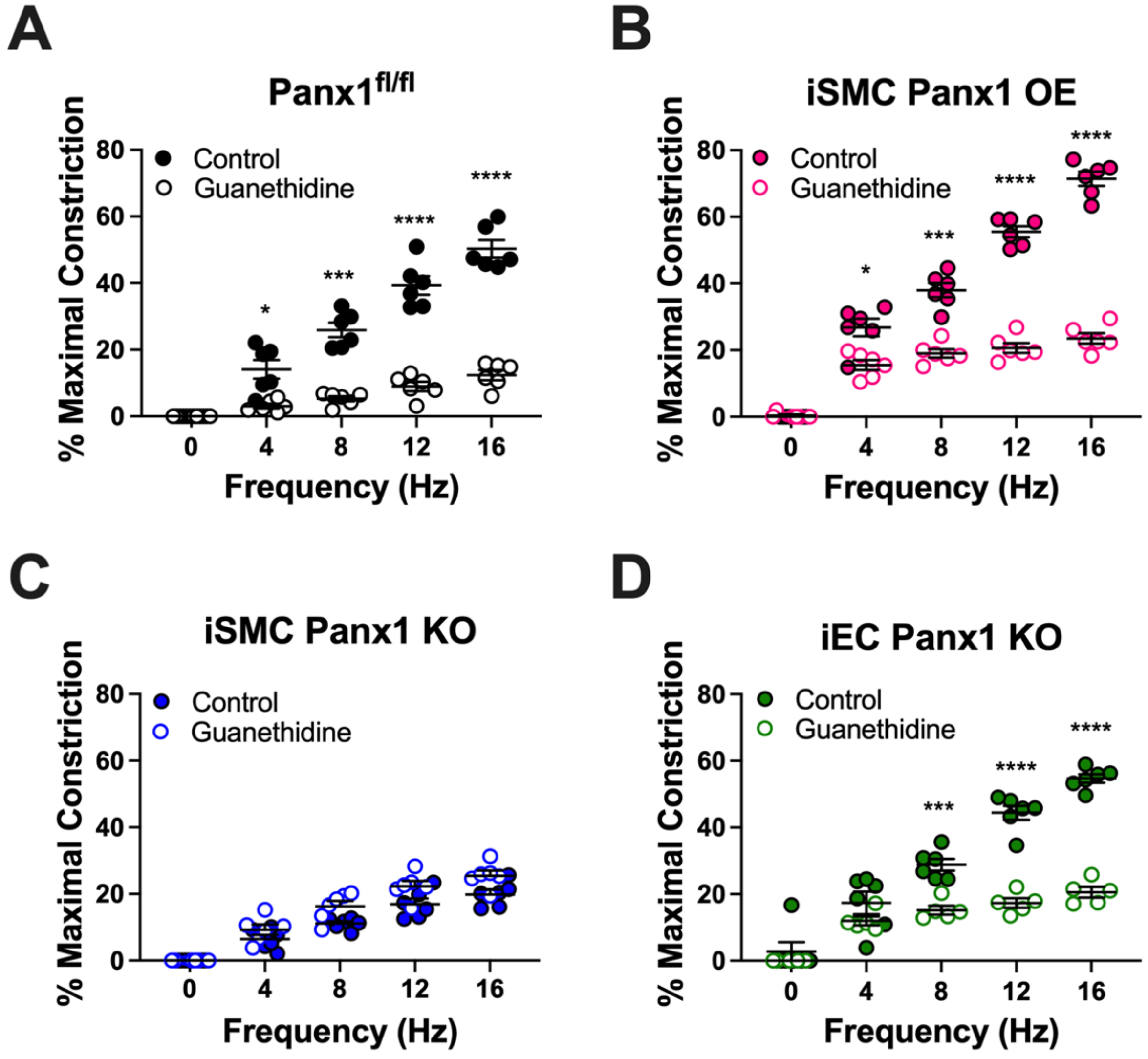


**Supplemental Figure 3: Disruption of norepinephrine release blunts sympathetic constriction.** (EFS)-induced constrictions with or without (A) guanethidine (10 μM) or (B) phentolamine (5 μM). Data were analyzed by repeated measures two-way ANOVA and Sidak post hoc test. * p<0.05, ***p<0.001, **** p<0.0001

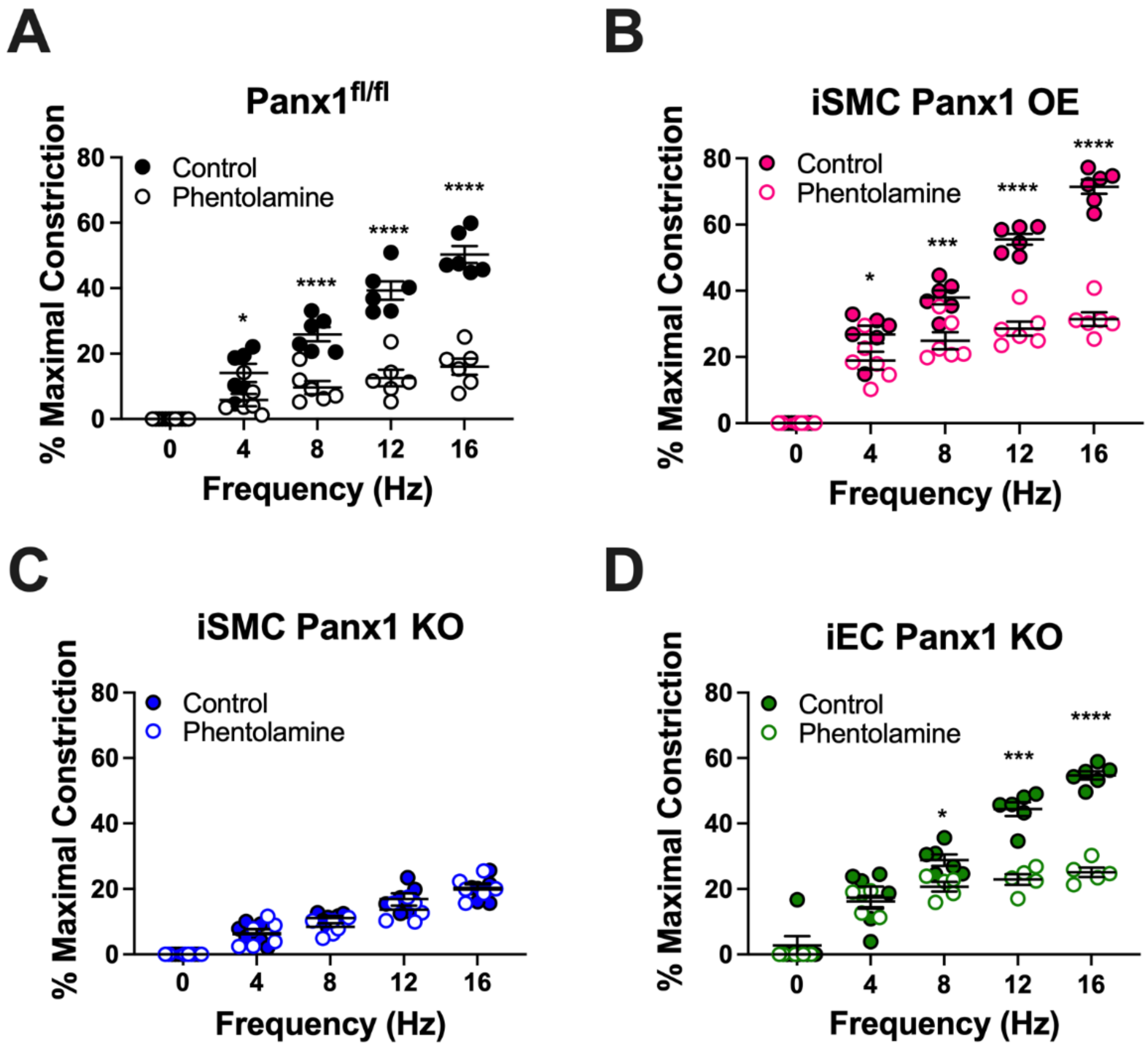


**Supplemental Figure 4: Alpha-adrenergic blockade blunts sympathetic vasoconstriction.** EFS-induced constrictions with or without phentolamine (1 μM). Data were analyzed by repeated measures two-way ANOVA and Sidak post hoc test. * $p<0.05$, ***$p<0.001$, **** $p<0.0001$

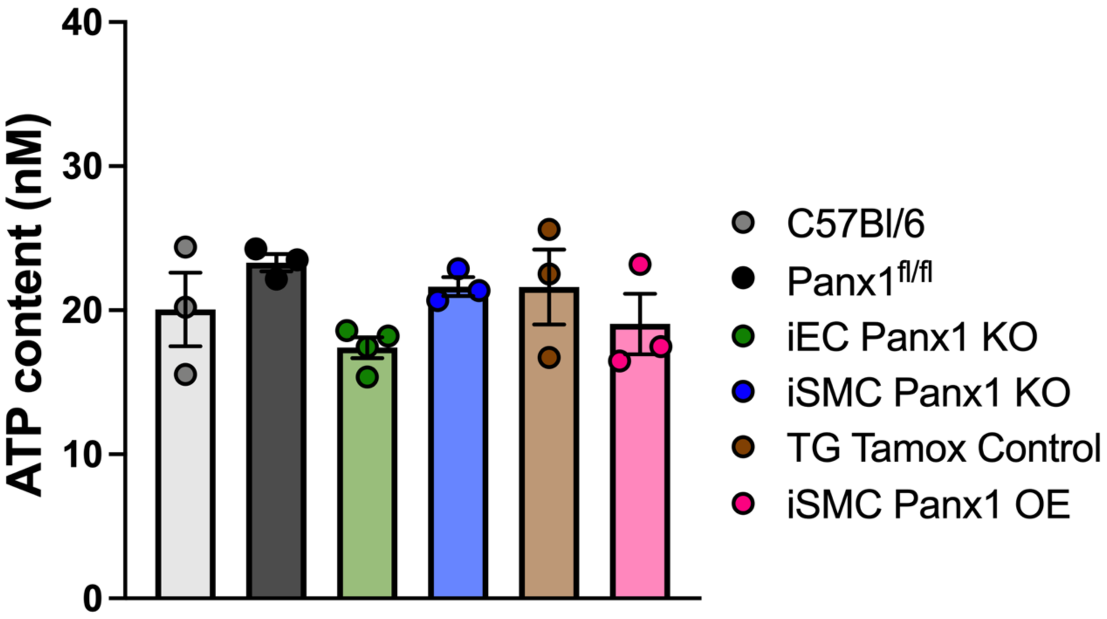


**Supplemental Figure 5: EC and SMC Panx1 does not change intracellular ATP.** ATP content of third order mesenteric arteries. Data were analyzed by one-way ANOVA and Holm-Sidak post hoc test.

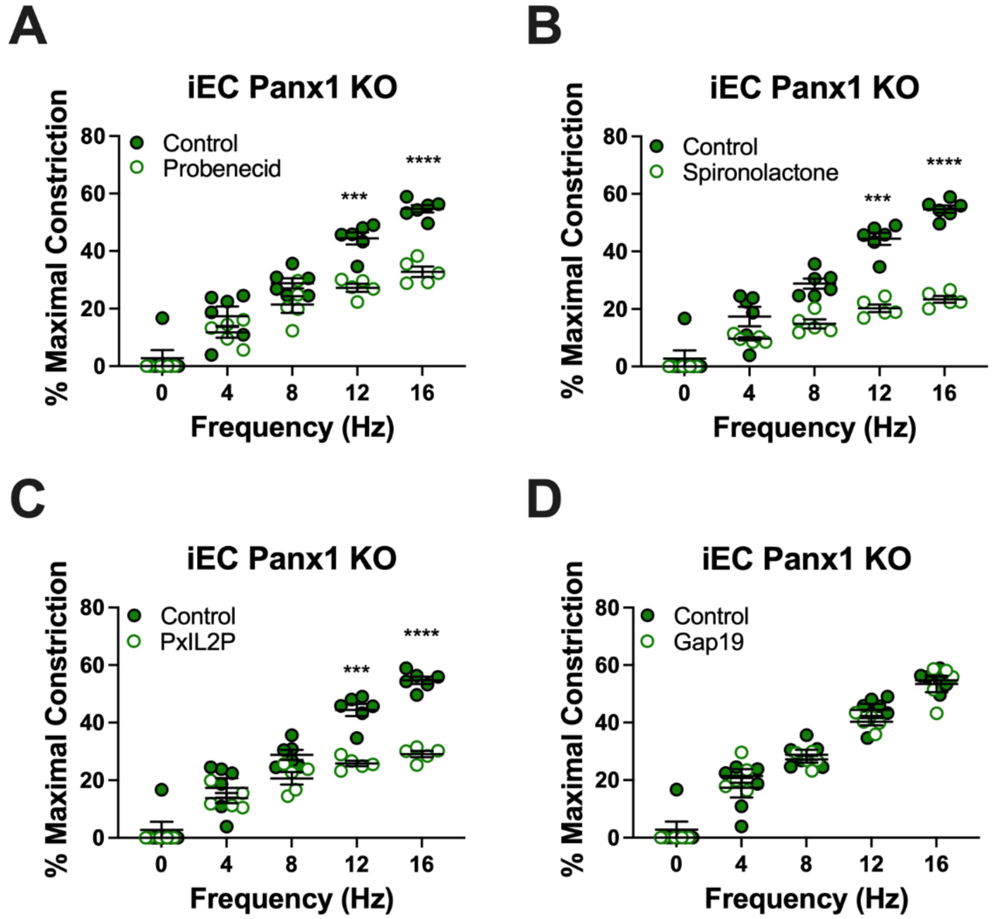


**Supplemental Figure 6: Panx1 inhibition blunts sympathetic vasoconstriction independent of endothelial Panx1 expression.** EFS-induced constrictions in iEC Panx1 KO mice with or without 2 mM probenecid, 20 μM spironolactone, 20 μM PxIL2P, and 20 μM Gap19. Data were analyzed by repeated measures two-way ANOVA and Sidak post hoc test. * p<0.05, ***p<0.001, **** p<0.0001

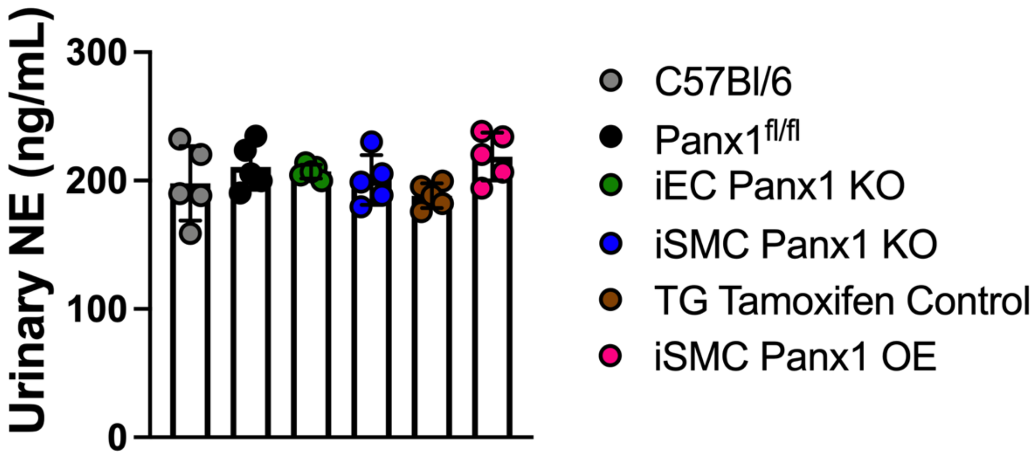


**Supplemental Figure 7: Urinary norepinephrine (NE) was not different between genotypes.** Data was analyzed by one-way ANOVA and Holm-Sidak post hoc test.
